# Insight into the genetic diversity, resistance, and virulence of *Listeria* from the marine environment: reveal the risk of hypervirulent isolates

**DOI:** 10.1101/2023.04.06.535972

**Authors:** Mao Pan, Wang Yan, Li Lingling, Ji Shunshi, Li Peijing, Liu Lingyun, Chen Jinni, Sun Hui, Luo Xia, Ye Changyun

## Abstract

*Listeria monocytogene*s is a major human foodborne pathogen and a ubiquitous environmental saprophyte. In this study, we investigated the prevalence and characteristics of *Listeria spp*. from beach sand in the coastal environment. Three different *Listeria spp*., *Listeria monocytogenes* (n=16), *Listeria fleishmanii* (n=7), and *Listeria aquatica* (n=3) were isolated from 769 beach sand samples and demonstrated diverse biofilm forming capacity. The *L. monocytogenes* isolates belonged to four distinct sequence types (ST87, ST121, ST35, and ST85) and contained the majority of virulence genes, some isolates were hypervirulent clones or had close phylogenetic relatedness with clinical cases. The ST87 isolates showed higher ability of biofilm formation in seawater than other STs strains. As a reservoir of microbes from marine environments and human/animal excrement, coastal sand would play an important role in the spread of *L. monocytogene*s and is an environmental risk for human listeriosis.

## Introduction

The genus *Listeria* comprised 30 species according to the List of Prokaryotic Names with Standing in Nomenclature (LPSN). *Listeria monocytogenes*, the main human pathogenic species, is an important opportunistic bacterium and the etiologic agent of listeriosis in elderly, pregnant women, children, and immunosuppressed patients, which can cause septicemia, meningitis, abortion, and even fatal infections (1). Additionally, *L. monocytogenes* is an environmental saprophyte that occupies a wide range of habitats, including river, soil, vegetation and food processing environments (1). *L. monocytogenes* is well adapted to grow under harsh environmental conditions, such as high salt concentration, pH 4.5, heavy metals, and / or freezing temperature. The bacterium could survive on dusty sand particulates (10 °C and 88% relative humidity) for more than 151 days (2). *Listeria fleishmanii* was initially isolated from cheese and has persisted for more than 5 years, and it has also been found sporadically in soil and feed (3, 4). *Listeria aquatica* was first identified in water in the United States (5). *L. monocytogenes* was reported to be constant presence in the lagoon and soil around the pig manure treatment plants (6). The species *L. monocytogenes* had regional heterogeneity, ecological niche preference, and genetic diversity, with four genetic lineages (I-IV) and 14 serotypes and were further subdivided by clonal complexes (CCs) and multi-locus sequence types (STs). The majority of clinical isolates were belonged to lineage I strain, while the food isolates were concentrated in lineage II (7). Serotype 4b strains were associated with human infections worldwide, whereas ST87 strains of serotype 1/2b were the predominant clinical clones in China (8, 9).

It has been well documented that infection of *L. monocytogenes* is derived from the consumption of contaminated food, such as ready-to-eat meat, soft cheese, and unpasteurized milk. Pathogenic marine bacteria-contaminated seafood was a threat to human health, particularly in coastal cities (10, 11). Water and soil are also sources of *L. monocytogenes* infection (12). An immunodeficiency adult was reported to have septic arthritis from *L. monocytogenes* after swimming in a public pool (13). Migratory wild birds, such as the black-headed gull, inhabit a variety of aquatic environments, which can facilitate the geographical spread of *L. monocytogenes* (14).

Environmental surveys of *L. monocytogenes* on the central California coast showed a prevalence of 41.9% to 62% in public access surface water with serotype 4b strains accounting for most of the isolates (15-17). Further analysis revealed 90% of *L. monocytogenes* isolates from central California Coast contain *inlA* gene encoding for intact InlA, which differs significantly from isolates found in foods and food processing environments (18). Another research of *L. monocytogenes* isolates in Morocco showed a prevalence of 2.5% in the marine environment with serotype 1/2b strains in the majority (19). The presence of *L. monocytogenes* in a variety of marine habitats would be the potential contamination source and a threat for human health.

To date, the epidemiological investigation and virulence assessment of *L. monocytogenes* in the marine environment in China have not yet been reported. Therefore, a survey for the prevalence and characteristics of *Listeria spp*. in beach sands was first explored in a coastal city in China.

## Results

### Occurrence and molecular subtyping of *Listeria* isolates from beach sand

A total of 26 isolates of three distinct *Listeria* species were obtained from 3.38% of 769 sand samples collected along the coast. The 16 *L. monocytogene*s strains belonging to two lineages (I and II) and four sequence types (ST87, ST121, ST35, ST85) were isolated from 2.08% of the samples in bays B, D, E, village V and island I. The prevalent ST87 strains were found in B, D bays, V village and I island. Seven strains of *L. fleischmannii* were found in B, C, and E bays with a total prevalence of 0.91% and prevailing in C bay. Three strains of *L. aquatica* was found sporadically in E bay with a total frequency of 0.39% (Figure 1 A, B).

**Figure 1.**
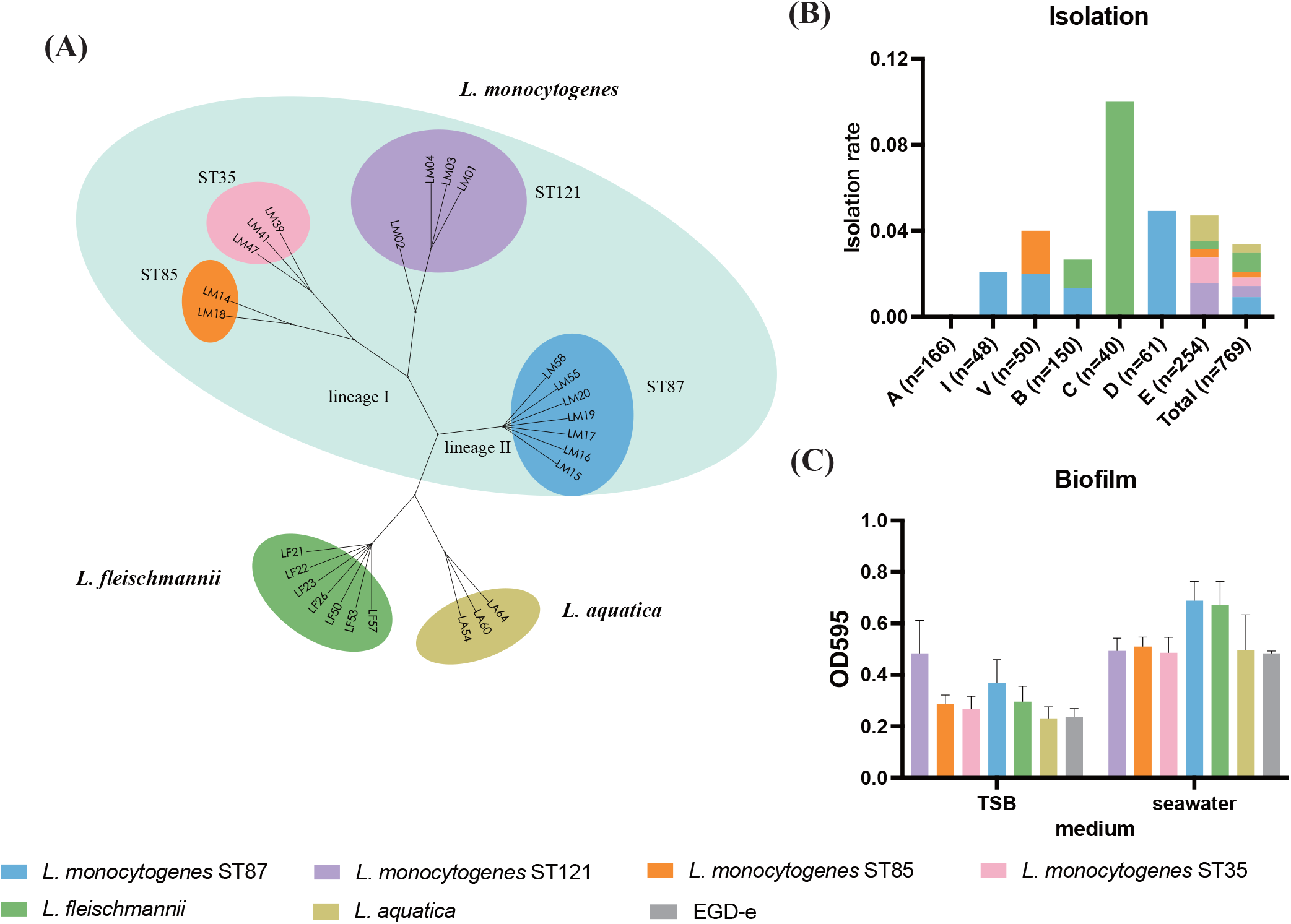
(A) Unrooted phylogenetic tree based on core genes of *Listeria* isolates collected in this study (n=26). (B) Isolation rate of *L. monocytogenes, L. fleischmannii*, and *L. aquatica* in each site. (C) Biofilm formation of *Listeria* in TSB and seawater.

### Virulence genes, resistance genes, and plasmid of *Listeria monocytogenes* isolates

A total of 182 virulence and resistance genes were screened among isolates in this study. All 16 *L. monocytogene*s isolates contained the majority of virulence genes and resistance genes (Figure 2). The LIP-1 (including *plc, actA, prfA, plcB, hly* and *mpl*), *inlA* (full-length), *inlB, inlC*, and *iap* were detected in all *L. monocytogene*s isolates from the beach sand. LIPI-3 was not found in these *L. monocytogene*s isolates, and LIPI-4 was found in ST87 isolates. The ST121 isolates contained stress survival islet 2 (SSI-2), while the ST35 and ST85 isolates contained stress survival islet 1 (SSI-1). The ST121 isolates carried cadmium resistance genes *cadA1C1*, while the ST35 isolates contained genes *cadA3C3*. Disinfectant resistance genes were not detected in these isolates. All *L. monocytogene*s isolates contained intrinsically antibiotic resistant genes *norB (*fluoroquinolones resistance*), fosX* (fosfomycin resistance) and *mprF* (cationic peptides resistance) according to CARD database.

**Figure 2.**
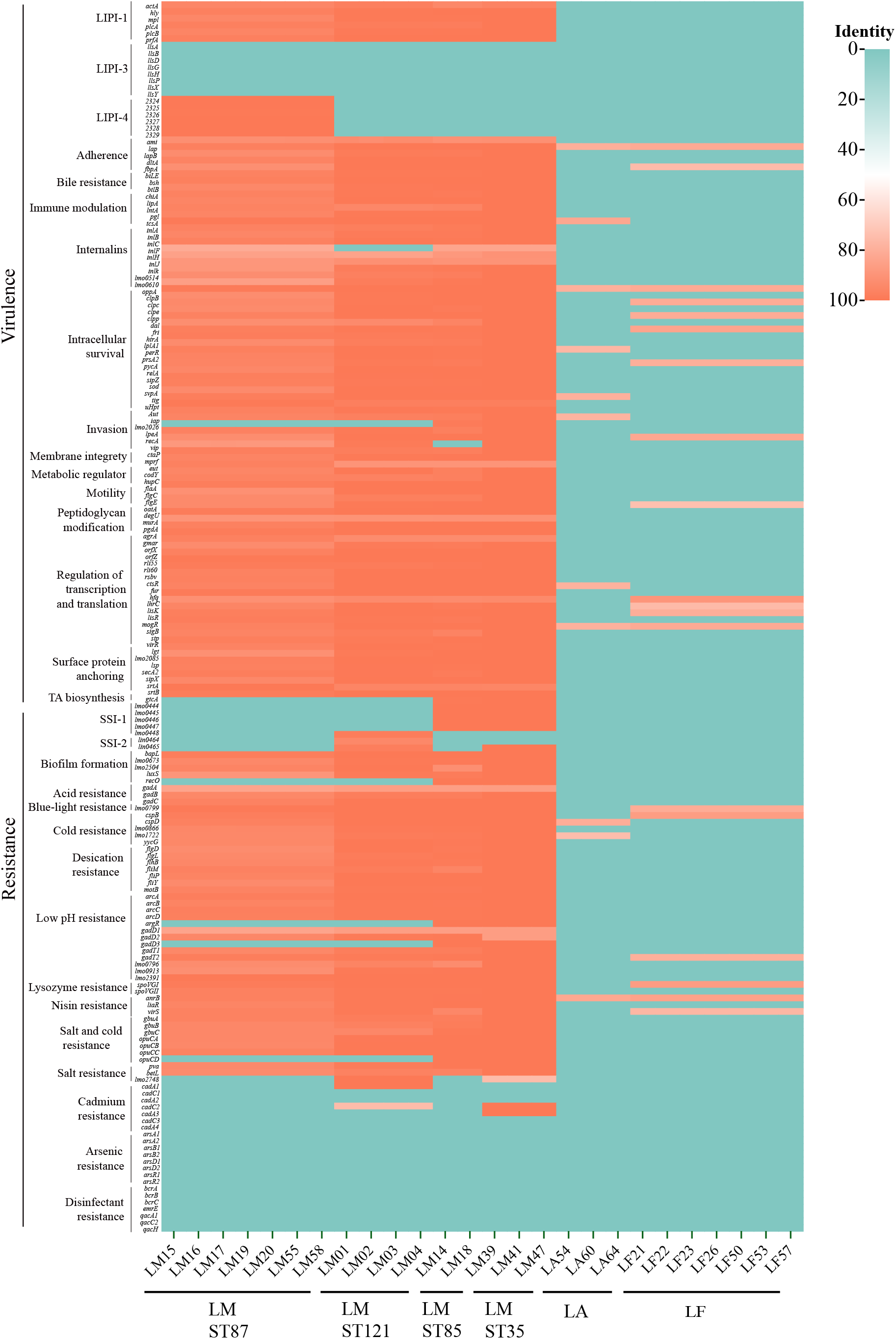
Color heatmap of presence (Green)/ absence (Orange) gene matrix with 187 virulence and resistance genes comprising the *Listeria* isolated in this study.

ST121 isolates of *L. monocytogene*s contained a 57 kb plasmid, which was found in different *Listeria spp*. strains in USA, Italy, Australia, Poland and Russia (Table 2). This plasmid was discovered in two human source isolates of ST7 and ST122 (20, 21). It contained 67 coding sequences (CDS), encoding ten transposase elements, and many functional proteins, such as putative UV-damage repair protein UvrX, type II toxin-antitoxin system, CRISPR-associated protein Cas5, cadmium-transporting ATPase, copper-translocating P-type ATPase CopB, glycine/betaine ABC transporter GbuC, NADH peroxidase Npx (Figure 3).

**Table 1.** The information of Listeria monocytogenes genomes used in this study.

| Country | ST121 | ST87 | ST35 | ST85 |
| --- | --- | --- | --- | --- |
| United Kingdom | 690 | 31 | 2 |  |
| USA | 261 | 276 | 21 | 4 |
| France | 89 |  |  |  |
| Canada | 42 | 24 | 1 |  |
| China | 42 | 260 | 6 | 2 |
| Switzerland | 24 |  |  |  |
| Australia | 24 | 2 | 1 | 1 |
| Italy | 22 | 1 |  |  |
| Ireland | 16 | 1 |  |  |
| Belgium | 15 | 5 |  |  |
| Norway | 13 |  |  |  |
| Spain | 12 | 17 |  |  |
| Germany | 12 | 17 |  |  |
| Netherlands | 11 | 9 |  |  |
| Poland | 7 | 1 |  |  |
| Russia | 6 |  |  |  |
| Denmark | 5 |  |  |  |
| Mali | 5 |  |  |  |
| Senegal | 4 |  |  |  |
| Latvia | 3 |  |  |  |
| Iceland | 3 |  |  |  |
| South Africa | 3 | 1 |  |  |
| Finland | 3 |  |  |  |
| Venezuela | 2 |  |  |  |
| Chile | 1 |  |  |  |
| Uruguay | 1 |  |  |  |
| Turkey | 1 |  |  |  |
| South Korea |  | 9 |  |  |
| Indonesia |  | 4 |  |  |
| New Zealand |  | 1 |  |  |
| Portugal |  | 1 |  |  |
| South Africa |  | 1 |  |  |
| Unknown |  |  | 1 | 3 |
| Total | 1317 | 661 | 32 | 10 |

**Table 2.**
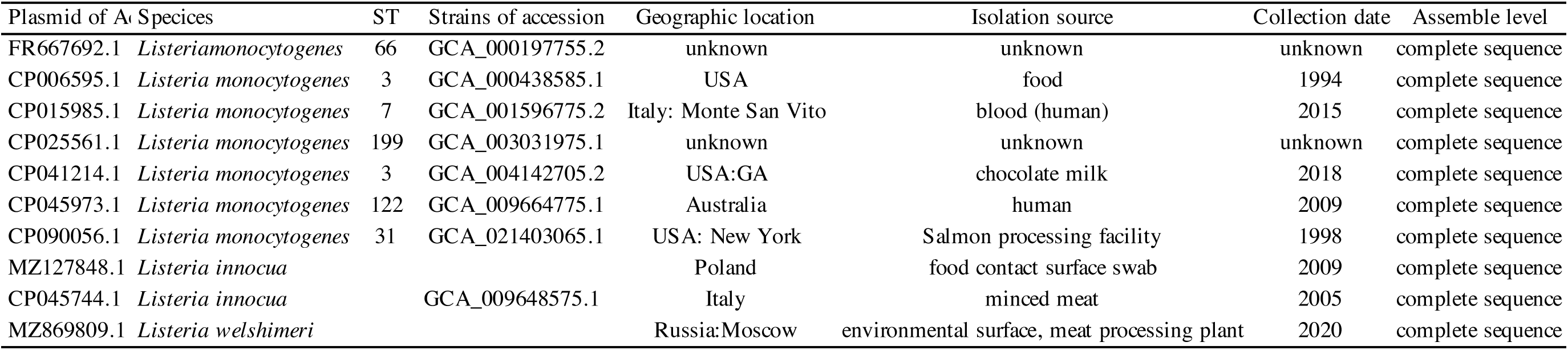
The global plasmid information.

**Figure 3.**
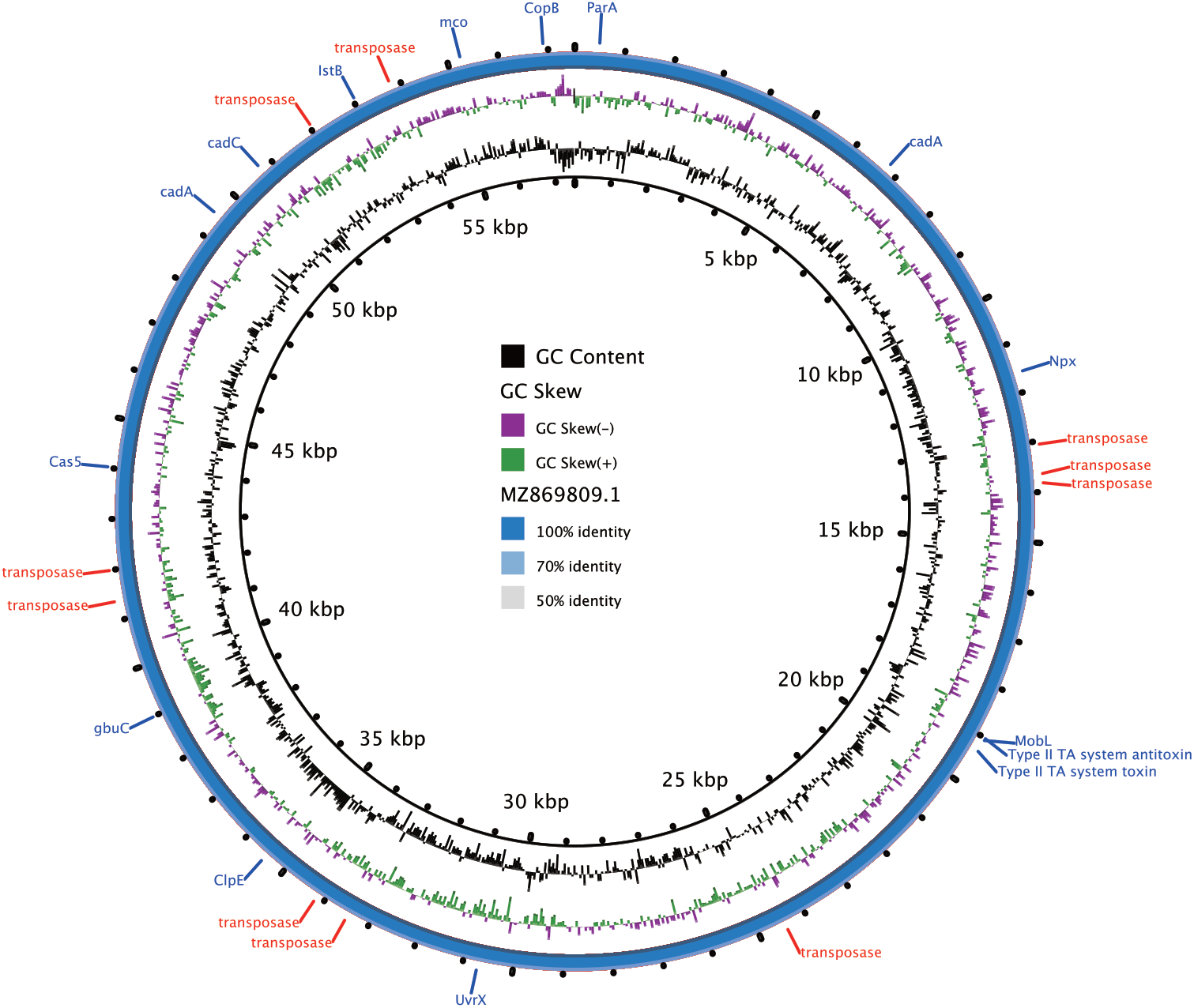
The plasmid pMZ869809.1 of complete sequencing as representative. BRIG was used to generate a visual representation of 57530-nt contig. GC contents and skew are presented in the inner rings; The annotation of this plasmid with known genes indicated is represented in the outer ring.

### Genetic relationship of *L. monocytogenes* isolates

Genome comparison was explored between seven ST87 isolates from beach sand and other 654 isolates of *L. monocytogenes* (264 isolates from human, 206 isolates from food and food processing environments, 95 isolates from environments, 11 isolates from animals and feed, and 78 isolates from unidentified sources). Phylogenetic analysis showed an obvious tendency of regional clustering, and the sandy isolates in this study had a closer-related evolution relationship with isolates in China than those in the United States. Isolates from clinical cases are distributed in the most evolutionary branch (Supplementary Figure 1). The complete LIPI-4 exists in all ST87 isolates (n=661), but not in other STs isolates (n=1358) (Supplementary Figure 5).

Four ST121 isolates of *L. monocytogenes* from beach sand in this study were compared with 1313 isolates from other regions (787 isolates from food and food processing environments, 375 isolates from environments, 122 from humans, 8 isolates from animals and feed, and 21 isolates from unidentified sources). The four sand isolates shared the closest relationship with four isolates from three clinical cases between 2007 and 2008 in China with 36-51 SNP differences (Supplementary Figure 2). In addition, ST121 (1307 isolates, 99.2%) contained complete SSI-2, whereas the other strains lack this island (Supplementary Figure 5).

Compared with 29 ST35 isolates from other sources, the three isolates from beach sand were closest to the three isolates from Dianchi Lake black-headed gulls in China with 1-8 SNP differences (Supplementary Figure 3).

For the ST85 strains, two isolates from beach sand and 8 isolates (5 isolates from food and food production environments and 3 isolates from unidentified sources) from USA, China, and Australia isolates were observed close phylogenetic relatedness and had 1-9 SNPs differences with each other (Supplementary Figure 4).

### General genomic features of *L. fleischmannii* and *L. aquatica* isolates

For the 16 isolates of *L. fleischmannii* (7 isolates from beach sand and 9 from other sources), the core genome (1659 genes) is 29.5% smaller than that of the *L. monocytogenes* (2354 genes), while the pan genome (6170 genes) is 40.8% larger than that of the *L. monocytogenes* (4383 genes) (22). *L. fleischmannii* genome is smaller in size but carries more accessory genes, which are associated with species diversity and confer competitive benefits on individuals. For the five isolates of *L. aquatica* (2 isolates from beach sand and 3 from other sources), the core genome (2063 genes) and pan genome (3851 genes) were 12.4% and 12.1% smaller than *L. monocytogenes* genome (Table 3) (22). The virulence genes in *L. fleischmannii* and *L. aquatica* were obviously less than those in *L. monocytogenes* (Figure 2). *L. fleischmannii* harbored fluoroquinolones resistance gene *norB*, while *L. aquatica* did not carry antibiotic-resistant genes. Plasmid was not found in *L. fleischmannii* and *L. aquatica* in this study.

**Table 3.**
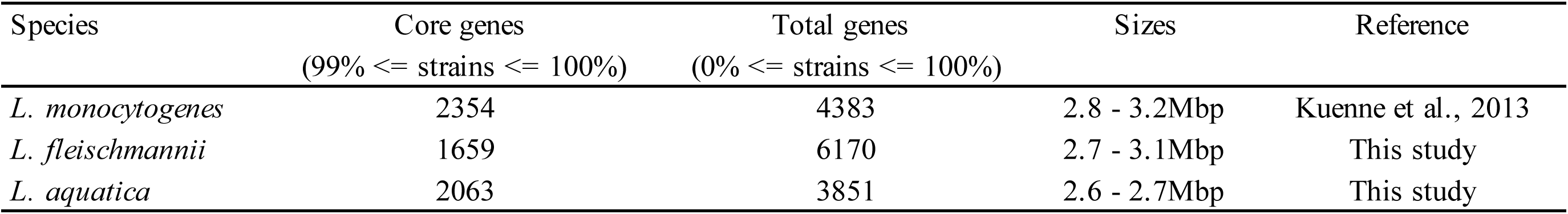
The core and pan genome information of *Listeria*.

### Biofilm formation ability of *Listeria* from beach sand

*Listeria* displayed stronger biofilm in seawater than TSB medium (Figure 1C). *L. monocytogenes* ST87 strains and *L. fleischmannii* isolates exhibited significantly stronger biofilm formation in seawater than isolates of other STs of *L. monocytogenes* and *L. aquatica*, while ST121 isolates exhibited significantly higher biofilm formation ability in TSB medium compared with other *Listeria* strains in this study.

All *L. monocytogenes* in this study harbored known biofilm-associated genes (*bapL, lmo0673, lmo2504, luxS, recO*) and its regulation gene (*argA, Rli60, sigB, mogR*), which were genetically polymorphic in different sequence types of isolates (Figure 2).

## Discussion

The cumulative annual cost of water-related illnesses in California was $3.3 million, which was caused by coastal water pollution for only two beaches (23). The occurrence of Cholera outbreaks were associated with aquatic environment in lakeside areas by using time-series analysis of 85565 cholera cases (24). *Escherichia coli* O157:H7, associated with haemorrhagic colitis and haemolytic uraemic syndrome, has been isolated in marine environments and could persist for up to a year in various water environment (25). *L. monocytogenes* is an important foodborne pathogen that is capable of infecting both human beings and animals. Multiple transmission modes besides food contamination have been reported to, such as raising or playing with animals, contacting with contaminated water and environment, etc (12, 26, 27). Recreational water activities on the coast polluted with fecal contamination were associated with gastrointestinal disease (> 120 million cases) and respiratory disease (> 50 million cases) each year around the world (28, 29). It is a potential threat to human health to exposure to beach sand and recreational seawater with *L. monocytogene*s contamination. To date, limited information was known about the prevalence and molecular characteristics of *L. monocytogene*s from beach sand, and no study on the prevalence and risk of *L. monocytogene*s in the marine environment from China was reported. The identification of *Listeria* genetic features in marine environment based on whole genome sequencing would be useful for risk assessment and traceability analysis.

In this pioneering study of *Listeria* from beach sands in a tourist city, we showed 26 out of 749 beach sand samples from the coastal environment were positive for *Listeria spp*. and revealed the existence of hypervirulent isolates (such as the ST87 strains) in marine environment. The sand would serve as an important environmental reservoir of *L. monocytogene*s with the specific biofilm formation ability.

ST87 was one of the main sequence types *L. monocytogene*s in raw seafood and fresh aquatic products in China (30, 31), and also the predominant sequence type in clinical infection in China and rarely reported in Europe and the United States (9, 32, 33). In this study, the ST87 isolates of *L. monocytogene*s were prevalent strains in beach sand and exhibited stronger biofilm formation than other isolates in seawater, which could contribute to the prevalence in aquatic products and cause the problem of food security. The ST87 isolates showed regional heterogenicity and the sandy isolates had closer relationship with those in China (over 150 isolates from human and 84 isolates from food and environments in China). Over 200 isolates (more than 150 isolates from food and environments and 46 isolates from human in USA) have emerged in the United States in NCBI database. Total of 661 ST87 isolates around the world contained the majority of virulence genes and LIPI-4, a cluster of six genes referred to a transplacental and central nervous system infection, which is consistent with a previous study about ST87 strain related sporadic listeriosis cases (32). The hypervirulent isolates ST87 presented in the marine environment might be a contamination source for seafood and a risk factor for public human health.

ST121 strains is highly abundant in food and was found to persist in food-associated environments (34). ST121 isolates in this study exhibited stronger ability than other isolates in nutrient-rich medium TSB, which might contribute to the prevalence and persistence in food and food-processing environments. The four ST121 beach sand isolates were grouped together with four human isolates from infant cord blood (2 isolates), placenta, and ascites (35). These eight isolates carried *inlA* gene encoding a full-length InlA with the same amino acid sequence and harbored a 57 kb multi-functional plasmid, which were different form other reported ST121 isolates(36). It was reported that 81.4% of ST121 isolates carried a conserved plasmid 62kb pLM6179, and 97.1% of isolates contain a truncated *inlA* gene (36). The ST121 strains infection was less reported in humans, the association between the ST121 isolates of sandy with the clinical strains would need further epidemiological investigation.

Plasmids in *L. monocytogene*s could benefit the survival and enhance the virulence of LM populations (37, 38). The 57kb plasmid was found in ST121 strains in this study, and existed in diverse host strains in many countries between 1994 and 2020, which codes many functional and uncharacterized proteins, including UvrX for resistance to UV-induced damage, GbuC for resistance to osmosis, Npx for resistance to oxidative stress, type II TA system for plasmid maintenance, and transposases facilitating genomic islands across diverse bacteria (38-40). This plasmid might contribute to the tolerance of *L. monocytogene*s against sunlight exposure and sea salt in beach environment.

Seabird can carry pathogenic microbes, and act as the long-distance disseminator of infectious agent though migration (14). A total of 2899 black-headed gulls occurred on this region between 2008 and 2020 in annual waterbird surveys (41). In this study, ST35 isolates in beach sand had closet relationship with the isolates from black-headed gulls in 2016, suggesting these isolates probably come from the same source, and presented long-distance spread by black-headed gulls migration. Fecal contamination with *L. monocytogene*s dropped in the sand might result in the persistence of the pathogen in the recreational marine environments and cause infection risk for people around the beach. Therefore, the potential risk of bird-mediated *L. monocytogene*s infection would not be ignored.

In the seawater and river environments, the bacterial community varied with the geographical location with greater species abundance in the downstream due to various factors, such as human activity, algal blooms (42, 43). In this study, bays A/B/C/D/E were located up and downstream of the sea in the coastal city, *Listeria spp*. were positive with diverse molecular characteristics in all sampled spots except bay A. Located in the upstream of these bays, bay A has the clearest coastal water and a low population density. Bay B is about 10 kilometers away the downtown area with white sand beaches. Bay C is a niche tourism destination and enriched with coral reef, which located approximately 4 kilometers away the downtown area and had the most isolates of *L. fleischmannii*. Bay D located in the downtown area and offers a public bathing beach with visa-free access, had a relatively high isolation rate of *L. monocytogenes*. Bay E, the downstream of these bays had a relatively high species diversity of *Listeria*. Island I and village V were leisure resort and located between bay A and bay B, providing maritime entertainments and sports for tourists. The prevalence of *L. monocytogene*s increased with the degree of human activity was also reported in the research of Danish aquatic and fish-processing environments (44), indicating the correlation between human activities and *L. monocytogene*s isolation.

Biofilms of marine bacteria are pervasive occurrence in marine ecosystem, which contribute to rapidly adaptation to environmental challenges and involve in a set of functions (45). All *Listeria* isolates in this study could generate biofilm in seawater, particularly *L. monocytogenes* ST87 and *L. fleishmanii* isolates showed high biofilm formation ability. The presence of resistance genes for adaptation to environment, especially salt resistance (*gbuABC, opuCABC*), UV-resistance (UvrX), would help *L. monocytogenes* survive and proliferate under the marine environment. *L. fleischmannii* might have some unidentified biofilm formation genes that need further study. The seawater contamination by *Listeria monocytogenes* signifies a constant risk of aquatic products, entering the food chain and posing a potential health risk for *Listeria* infection in humans. It is well known that the seafood contamination of *L. monocytogenes* emerged continuously, and 0.5-1.0 % of listeriosis cases were associated with ready-to-eat seafood (46). In addition, water recreational activities also have the exposure risk for people to infect *L. monocytogene*s by accidental ingestion.

Beach sands, as reservoirs of bacterial pathogens, were found for 10-to-100 fold higher bacterial abundance than adjacent recreational waters (47), and acts as an accumulation substrate and pollution source of bacteria from diverse sources, such as sewage disposal, and fecal dropping from wild birds and recreational users. However, there is little published data on the presence of *L. monocytogenes* in marine-associated environment and its risk level to human infection. We demonstrate that *Listeria* can produce biofilm in marine-associated environment, implying the possibility of persistence in the seawater and beach sand, which would cause the seafood contamination and infection of human. The susceptible individuals (eg, the immunocompromised or elderly people) were advised not to engage in beach recreational activities or expose their wounds to the marine-associated environment. Monitoring the pathogenic bacteria including *L. monocytogenes* by whole-genome sequencing and the fecal indicator bacteria level in the sands of recreational beaches is important for modeling public health hazards and environmental protection in the coastal cities.

Our study reveals the genetic and phenotypic characteristics of different species and sequence type isolates of *Listeria* in the coastal environment of China, showing the potential risk of *L. monocytogenes* infection in the marine environment. We found that the hypervirulent ST87 *L. monocytogenes* is the predominant sequence type in recreational sand beaches and had a high capacity for biofilm formation in seawater. The discovery of disease-associated sequence types of *L. monocytogene*s in the beach sand suggests the marine environment is an important reservoir for high-virulent *L. monocytogene*s. A routine surveillance system for pathogens in highly populated coastal regions is necessary for the public health.

## Materials and Methods

### Sand sampling and isolation of *Listeria spp*

A total of 769 sand samples were collected from five bays (A/B/C/D/E from north to south), an island, and a village on the path to the island in a coastal city in China, all sampling spots were accessible places for locals and tourists. Approximately 200g of sand from each sample was stored in sterilized plastic bags. Before the isolation of bacteria, the sand was socked and rubbed in approximately 100 ml of PBS. After the natural deposition of sand, the supernatant was transferred into a 50 mL tube for centrifugation at 4000 rpm for 10 min. The pellets were then spiked into half-Fraser and Fraser broth (Oxoid) for the enrichment of *L. monocytogenes* (48). The enriched culture was then streaked on Brilliance Listeria Agar (Oxoid) and incubated at 37 °C for 48 h. The presumptive positive colonies were selected for PCR identification using primers for *Listeria* genus-specific gene *prs* (Forward: *GCTGAAGAGATTGCGAAAGAA*; Reverse: *CAAAGAAACCTTGGATTTGCGG*).

### DNA extraction, Sequencing, Assembly, and Species identification

The *Listeria spps*. isolates obtained from sand samples were cultured in BHI at 37°C. Genomic DNA was extracted using the Wizard® Genomic DNA Purification Kit and quantified using the Qubit 2.0 Fluorometer. The DNA library was performed with Illumina using the NEBNext® UltraTM DNA Library Prep Kit. Whole genome shotgun sequencing of these isolates was performed on the Illumina Hiseq PE150 platform. High-quality paired reads were assembled using the SOAP denovo and SKESA (v 2.1). For species-level identification, fastANI (v 1.33) was used calculate the Average Nucleotide Identity (ANI) between genomes. Genome annotations were performed using Prokka (v 1.14.6).

### Core and pan genome and SNP-based phylogenetic analyses

WGS was used for the molecular characterization analysis of *Listeria spp*. The core and pan genome were analyzed with Roary pipeline using a 95% identity cut-off. The phylogenetic tree based on the core genes of *Listeria spp*. was constructed using FastTree. Sequence types were determined with *in silico* MLST. The publicly available strains from the NCBI database sharing the same sequence type with the sequenced isolates in this study were chosen for SNP analysis (Table S1). The geographic location of each ST strains was summarized in Table 1. The SNP alignment was performed using Snippy (v 4.6.0) and the phylogenetic tree was produced with Gubbins (v 2.4.1). The resulting phylogeny was plotted and visualized in ChiPlot (https://www.chiplot.online/) tools.

### Virulence genes, resistance genes, and plasmid analyses

Virulence- and resistance-associated genes were identified by sequence alignment with the existing *Listeria* virulence database in VirulenceFinder (https://bitbucket.org/genomicepidemiology/virulencefinder_db/src/master/) and reported resistance genes (34, 49). *In silico* antibiograms and plasmids were identified using ABRicate (v 1.0.1) with CARD and plasmidFinder. The plasmid contigs were extracted using SPAdes (v 3.14.4) in the plasmidSPAdes mode.

### Biofilm formation

Biofilm formation assays were performed using the classical crystal violet method (50). 10^7^ CFUs of microbial suspensions and 200 μl of TSB / sterile seawater were transferred to a 96-well microtiter plate and incubated for 4 days at 25 °C. Suspended bacteria were rinsed three times with PBS. Cell layers were stained with 1% crystal violet solution for 30 minutes, and then washed with deionized water. The crystal violet bound to biofilm was dissolved in 95% ethanol and quantified at A595 by microplate reader (BioTek).

## Figure legends

Supplementary Figure 1 Whole-genome phylogeny of ST87 *L. monocytogenes* based on SNP. From outer to inner, the circles represent geographic location, isolation source, collection date, and LIPI-4 matrix.

Supplementary Figure 2 Whole-genome phylogeny of ST121 *L. monocytogenes* based on SNP. From outer to inner, the circles represent geographic location, isolation source, collection date and SSI-2 matrix.

Supplementary Figure 3 Whole-genome phylogeny of ST35 *L. monocytogenes* based on SNP. The geographic location and isolation source is presented in circular and square of different color block.

Supplementary Figure 4 Whole-genome phylogeny of ST85 *L. monocytogenes* based on SNP. The geographic location and isolation source is presented in circular and square of different color block.

Supplementary Figure 5 Color heatmap of presence (blue)/ absence (white) gene matrix with 187 virulence and resistance genes comprising the *Listeria* sequencing downloaded in NCBI.

